# Single-cell analysis of the human retina reveals stage-linked microglial states and neural-immune circuit rewiring in diabetic retinopathy

**DOI:** 10.1101/2025.02.07.637193

**Authors:** Luning Yang, Sen Lin, Yiwen Tao, Qi Pan, Tengda Cai, Yunyan Ye, Jianhui Liu, Yang Zhou, Quanyong Yi, Zen Huat Lu, Lie Chen, Gareth McKay, Richard Rankin, Yongqing Shao, Weihua Meng

## Abstract

Diabetic retinopathy (DR) is a major cause of vision loss worldwide. Here, we conduct single-cell RNA sequencing of twenty human retina samples (from living and post-mortem donors) across non-diabetic, diabetic, and DR states to create a comprehensive transcriptomic atlas. We identify two stable microglial populations—homeostatic and inflammatory—that exist along a functional continuum, plus a neutrophil cluster within C1QA+ myeloid cells with dynamic transitions occurring throughout disease progression. Module-level analysis reveals divergent transcriptional trajectories: homeostatic microglia maintain energetic programs while selectively upregulating stress elements, whereas inflammatory microglia layer additional pro-inflammatory programs onto preserved biosynthetic foundations. Eleven co-expression modules organize into two major axes: an inflammatory-stress axis, and a regulatory/metabolic-motility axis, with a stable translation module persisting across disease stages. Cell communication analysis further highlights sophisticated neural-immune interactions, particularly between photoreceptors and microglia. Our findings provide insights into the complex cellular dynamics of DR progression and suggest potential therapeutic targets for early intervention.

## Introduction

Diabetic retinopathy (DR) is the leading cause of vision impairment and blindness among working-age adults globally, posing a significant public health issue. It is characterized by progressive neurovascular dysfunction, involving both neurodegeneration and microvascular changes [1]. The global number of adults with diabetes has surpassed 800 million, with its prevalence rising from 7% to 14% between 1990 to 2020 [2]. Among them, 22.27% are affected by DR, and the global age-standardized prevalence of blindness caused by diabetic eye disease has increased by 14.9% over the same period [3, 4]. This trend poses a significant and growing burden on global health. While significant progress has been made in the treatment of DR, substantial limitations remain. Anti-VEGF therapy, panretinal photocoagulation, and corticosteroid treatments have demonstrated efficacy, but they are associated with incomplete responses, visual side effects, and complications [5, 6]. These therapies primarily target downstream effects rather than addressing the underlying pathogenic mechanisms [7].

The key gap in DR treatment is that current therapies focus on late-stage manifestations, with insufficient intervention in early molecular events. As a result, the complex cellular and molecular mechanisms driving disease progression are not adequately addressed [8]. Therefore, a comprehensive understanding of pathophysiology is needed to enable intervention in the early stages when the disease is more controllable. Targeted treatment at the cellular level, rather than addressing downstream complications, is crucial. This requires a deeper understanding of single-cell and molecular mechanisms [9].

While single-cell RNA sequencing (scRNA-seq) has emerged as a powerful tool for understanding complex biological systems, its application in DR research faces several critical limitations. The major challenge in the application of scRNA-seq in DR research is the difficulty in obtaining human retina DR samples. While a few single-cell transcriptomic studies have generated retinal atlases, most published datasets utilize healthy donor eyes rather than samples from specific disease conditions, limiting our understanding of pathological states. To date, while several studies have examined retinal transcriptomes in animal models or healthy human retinas, single-cell analysis of complete human DR retina samples remains notably absent. Two recent studies focused on fibrovascular membranes, surgically harvested vitreous and peripheral blood from proliferative diabetic retinopathy (PDR) patients [10, 11], but neither fully recapitulates the complete pathogenesis of human DR as they focused on specific tissue types rather than the retina. Another challenge is the rapid post-mortem degradation of retinal tissue. In addition, the high cost of scRNA-seq and the difficulty in obtaining human retinal tissues hinder its large-scale application in DR and phased experimental design.

To investigate DR in retinal samples, more and more recent studies have focused on microglial cells, as these resident tissue macrophages play a pivotal role in initiating and exacerbating retinal inflammatory responses [12]. Our study aims to address these fundamental limitations by utilizing twenty human retinal samples spanning three disease stages, collected from both living donors and post-mortem specimens with minimal processing time to maximize transcriptional fidelity. We employed an integrated analytical approach combining cellular characterization, modular co-expression analysis, and intercellular communication mapping to comprehensively characterize the cellular and molecular features of DR progression.

Our integrated analysis reveals three critical insights that address fundamental gaps in the field. First, we demonstrate that retinal microglia exist along a functional continuum with two stable populations (homeostatic and inflammatory) that shift transcriptionally during DR progression through coordinated modular programs rather than wholesale phenotypic switching. Second, we uncover systematic remodeling of intercellular communication networks, particularly the pathological shift from CD44-mediated to CXCR4-mediated MIF-CD74 signaling. Third, we validate the persistence of functionally distinct rod photoreceptor subtypes across disease stages [13], each engaging specialized signaling repertoires that differentially shape retinal immune dynamics. This study represents one of the first comprehensive single-cell analyses of human retina DR progression, revealing sophisticated neural-immune interactions that could advance our understanding of DR pathogenesis and open new avenues for targeted therapeutic interventions.

## Materials and Methods

### Human donor eyes collections for scRNA-seq analysis

From June 2023 to February 2025, we obtained twenty retina samples from non-diabetic, diabetic, and DR patients under protocols approved by the ethics committees of the University of Nottingham Ningbo China, the Lihuili Hospital, and the Ningbo Eye Hospital. Two samples of living donors were acquired through surgical enucleation at the Lihuili Hospital. The remaining eighteen samples came from the Ningbo Eye Hospital, comprising seven paired eyes and one single eye from deceased donors and three surgical specimens from enucleated patients. We utilized complete retinal specimens rather than selective punch biopsies, enabling comprehensive analysis of the entire retinal structure from center to periphery. In this study, ’fresh retinal samples’ refers to two categories, the first is the living donor samples obtained during surgical enucleation and processed immediately within 10 min, and the second is post-mortem specimens processed within 6 h of death with strict temperature control without frozen. Table 1 provides detailed information on each sample.

**Table 1.** Retinal sample information.

| Sample | Donor ID | Tissue Source | Eye | Sex | Age(y) | Diabetic Status* | Eye Removal/Death Reason | Hospital† |
| --- | --- | --- | --- | --- | --- | --- | --- | --- |
| 1 | D1 | Live | Left | Male | 76 | DM | Corneal staphyloma | L.H. |
| 2 | D2 | Live | Left | Male | 55 | nonDM | Phthisis bulbi | L.H. |
| 3 | D3 | Live | Right | Male | 71 | nonDM | Corneal ulcer | Y.H. |
| 4 | D4 | Live | Left | Female | 50 | nonDM | Phthisis bulbi | Y.H. |
| 5 | D5 | Live | Right | Female | 68 | nonDM | Corneal perforation | Y.H. |
| 6 | D6 | Cadaver | Left | Male | 34 | nonDM | Cerebral hemorrhage | Y.H. |
| 7 | D7 | Cadaver | Left | Female | 92 | nonDM | Natural death | Y.H. |
| 8 | D7 | Cadaver | Right | Female | 92 | nonDM | Natural death | Y.H. |
| 9 | D8 | Cadaver | Left | Female | 55 | DM | Traumatic brain injury | Y.H. |
| 10 | D8 | Cadaver | Right | Female | 55 | DM | Traumatic brain injury | Y.H. |
| 11 | D9 | Cadaver | Left | Male | 55 | DR | Brainstem hemorrhage | Y.H. |
| 12 | D9 | Cadaver | Right | Male | 55 | DR | Brainstem hemorrhage | Y.H. |
| 13 | D10 | Cadaver | Left | Male | 35 | DR | Traumatic shock | Y.H. |
| 14 | D10 | Cadaver | Right | Male | 35 | DR | Traumatic shock | Y.H. |
| 15 | D11 | Cadaver | Left | Male | 84 | DR | Prostate cancer | Y.H. |
| 16 | D11 | Cadaver | Right | Male | 84 | DR | Prostate cancer | Y.H. |
| 17 | D12 | Cadaver | Left | Male | 69 | DR | Lung cancer | Y.H. |
| 18 | D12 | Cadaver | Right | Male | 69 | DR | Lung cancer | Y.H. |
| 19 | D13 | Cadaver | Left | Female | 87 | DM | Liver cancer | Y.H. |
| 20 | D13 | Cadaver | Right | Female | 87 | DM | Liver cancer | Y.H. |

\*DM, diabetes mellitus without diabetic retinopathy DR; diabetic retinopathy; nonDM, non-diabetic retinopathy nor diabetic. Because the tissues were obtained intra-operatively or post-mortem and detailed fundus imaging records were unavailable, the exact stage or subtype of diabetic retinopathy (e.g., NPDR vs PDR) could not be ascertained for the DR cases.
†L.H., The Ningbo Lihuili Hospital; Y.H., The Ningbo Eye Hospital.

### Single-cell RNA sequencing

Human retina samples were dissociated for scRNA-seq using the LK003150-1bx Dissociation Kit (Worthington Biochemical Corporation) following the manufacturer’s protocol. Tissue was placed on ice in a petri dish and manually cut into 2-4 mm pieces. After incubation in RPMI/enzyme mix (Miltenyi Biotec), the tissue fragments were transferred to a gentleMACS C tube for dissociation. After dissociation, the suspension was passed through a 70 µm cell strainer and centrifuged at 300 × g for 10 min at 4°C. The supernatant was removed, and 10 times volume of 1 x Red Blood Cell Lysis Solution (130-094-183, Miltenyi Biotec) were added to lyse red blood cells at 4°C for 10 min. Subsequently, 10 mL of 1x PBS containing 0.04% BSA was added to the suspension, which was then centrifuged again at 300 × g for 10 min. The suspension was mixed with AO/PI fluorescent dye (1:1 ratio) and incubated for 30 s, then cell viability was assessed for CountStar analysis. Viable cell suspension was resuspended in 1x PBS with 0.04% BSA to 700-1200 cells/µL (viability > 85%) for loading onto the 10x Genomics Chromium platform.

scRNA-seq libraries were made from 10,000 cells per sample using the Chromium Single Cell 3’ Library and Gel Bead Kit (10X Genomics). 150 bp pair-ended sequencing was performed on Illumina Novaseq platform. Raw sequencing data quality was assessed using FastQC to ensure that (1) no more than three ambiguous “N” bases in total were present; (2) less than 20% of bases had a Phred quality score lower than 5; and (3) residue adapter sequences were removed. The resulting sequencing outputs were processed using the 10X Genomics Cell Ranger V8.0.1 pipeline with default settings. Transcript reads were aligned to the 10x Genomics Human reference index (GRCh38-2020-A). Each cell was assigned a unique, sample-specific identifier, and feature-barcode matrices were generated using the chromium cellular barcodes.

### Quality control and doublets removal

For each sample, feature-barcode matrices were processed using the 10X Genomics Cell Ranger pipeline (https://www.10xgenomics.com/support/software/cell-ranger). We performed quality control and then doublet removal for each dataset to eliminate low-quality cells. Doublets were identified and removed using DoubletFinder (https://github.com/chris-mcginnis-ucsf/DoubletFinder). High-quality cells were selected based on the following criteria in the Seurat R package (v5.1.0) [14]: cells with a minimum gene count of 200, a UMI count greater than 400, and a mitochondrial read fraction of less than 20%. Cells passing these filters were retained for downstream analysis.

### Data integration and clustering

After initial processing, the twenty datasets were integrated by Scanpy (v1.10.2) [15]. We used scVI-tools (v1.1.2) [16] to perform dimensionality reduction and batch-effect correction. A k-nearest neighbor (KNN) graph was built from the resulting latent space, and Leiden clustering was applied at multiple resolutions to identify and refine retinal cell types. Previously reported marker genes’ expression of each cluster was used to annotate their corresponding retina cell type. This annotation workflow was further repeated to refine an accurate cell type annotation for the sub populations within each cell type. Sample bias introduced by the inclusion of fresh retinas from living donors was evaluated by analyzing the proportions of various cell types across all datasets.

### Cell disturbance analysis

Transcriptomic variations across nonDM (samples without diabetes mellitus), DM (samples with diabetes mellitus), and DR (samples with diabetic retinopathy) conditions were assessed using Augur (v1.0.3) [17] and odds ratio (OR) analysis. To avoid instability from rare or small populations and non-ubiquitous clusters, T cells, retinal pigment epithelium, neutrophils, and retinal ganglion cells were excluded, and the comparison was restricted to nonDM versus DR. This pairwise comparison was prioritized because Augur is designed for binary classification tasks and because nonDM versus DR exhibits the most pronounced transcriptomic remodeling, whereas DM exhibited minimal molecular alterations relative to nonDM. Augur calculated Area Under the Curve (AUC) scores to evaluate the predictive power of gene expression profiles for distinguishing nonDM from DR within each cell type, where higher AUC indicates stronger disease-associated perturbation. Label-permutation controls were performed to confirm that observed AUC values reflected true condition-expression associations rather than modeling artifacts. For compositional differences, we constructed a 2×2 contingency table with rows = that condition vs. all other conditions and columns = that cell type vs. all other cell types for each cell type and for each condition column (nonDM, DR), We computed the odds ratio and exact p value using Fisher’s exact test, then applied Benjamini-Hochberg correction. OR > 1 denotes enrichment in the column’s condition. This composition-aware metric complements Augur’s separability, helping distinguish proportional shifts from transcriptional perturbations.

### Differential expression and enrichment analysis

We performed differential expression (DE) to identify cell type or condition-specific markers using Seurat’s FindAllMarkers and FindMarkers functions with an FDR < 0.05 and |log2FC| > 1 threshold for significance. To robustly define cluster-specific marker genes, particularly for microglial subpopulations, COSG (v0.9.0) [18] was applied following sub-clustering of C1QA-positive myeloid cells, which prioritizes genes with high specificity and robustness for each annotated cluster. Gene Ontology (GO) and Kyoto Encyclopedia of Genes and Genomes (KEGG) enrichment analysis were conducted by clusterProfiler (v4.10.1) [19]. Pathway scoring and enrichment were further evaluated by the irGSEA (v3.3.2) [20] framework, combining multiple methods (AUCell, UCell, singscore, ssgsea, JASMINE, viper) through robust rank aggregation (RRA).

### Transcriptional pattern discovery

Microglia-specific gene co-expression modules were identified using Hotspot. First, Hotspot analysis (v1.1.1) identified spatially coordinated gene modules based on local correlation (https://github.com/YosefLab/Hotspot), leveraging dimensionality reduction and KNN graph construction. Module relationships were evaluated through correlation analysis, with scores calculated using UCell (v2.6.2) [21].

### Cell communication analysis

CellChat (v1.6.1) was applied to perform cell-cell communication analysis [22]. First, CellChat objects were created for DR and nonDM conditions using the human CellChatDB database. Overexpressed genes and interactions were identified with default parameters and the analysis was enhanced by projecting data onto protein-protein interaction networks. Communication probabilities were computed using the triMean method. Signaling pathway analysis was performed by aggregating the communication probabilities of all ligand-receptor pairs associated with each pathway. To compare communication changes between nonDM and DR conditions, we specifically examined how each microglial subtype and rod photoreceptor subtype acted as signal senders or receivers with respect to other cell types

## Results

### Construction of the human retinal scRNA-seq atlas and lineages perturbed across DR

To investigate diabetic retinopathy progression, we performed scRNA-seq on twenty human retina samples encompassing three disease stages (nonDM, DM, DR). After stringent quality control and data integration using scVI, the retinal cell populations were visualized in a uniform manifold approximation and projection (UMAP) plot (Figure 1A). Cells including cone, rod, bipolar (BC), amacrine (AC), horizontal (HC), retinal ganglion (RGC), Müller glia (MGC), astrocytes, microglia, T cells (TC), retinal pigment epithelium (RPE), and neutrophil, were identified according their significantly expressed distinctive marker genes (Figure 1B). Across donors, rods dominated in 19/20 samples, with group means markedly higher than any other lineage (nonDM 67.9 ± 18.1%, DM 56.5 ± 18.8%, DR 76.8 ± 6.3%; Table S1). For non-rod lineages, between-group differences were modest relative to within-group dispersion and showed substantial overlap. For example, MGC medians [IQR] were 10.8% [8.5–13.5%] in nonDM, 13.6% [13.3–30.5%] in DM, and 7.4% [6.2–9.9%] in DR; microglia were 1.8% [1.4–6.0%], 1.5% [0.4–2.6%], and 0.6% [0.5–0.7%], respectively. These distributions indicate notable inter-donor variability; accordingly, downstream analyses emphasize donor-level statistics and pair composition with within-lineage transcriptomic separability to distinguish proportional shifts from transcriptional perturbations. Then, we quantified disease-associated changes using a composition-aware metric (OR) and a perturbation-ranking approach (Augur) to explore a wider array of cell types associated with DR. To avoid instability from rare/small populations and a non-ubiquitous TC cluster, TC, RPE, Neutrophils, and RGCs were excluded, and the comparison was restricted to nonDM versus DR. Compositionally, the OR heatmap showed condition-specific enrichment among the remaining major lineages, for example, microglia were relatively enriched in nonDM, whereas cones appeared relatively enriched in DR (Figure 1C). In contrast, Augur prioritizes transcriptomic separability within each lineage, where higher AUC indicates stronger DR perturbation and permutation collapses values toward chance (≈0.5) (Figure 1D). The standard Augur run ranked HC highest (AUC = 0.682), followed by Cone (0.651), Microglia (0.649), Rod (0.625), and BC (0.593), with MGC, AST, and AC closest to chance. A label-permutation control yielded AUCs near 0.5 for all types, supporting that the observed ordering reflects true label–expression structure rather than modeling artifacts (Figure 1E).

**Figure 1.**
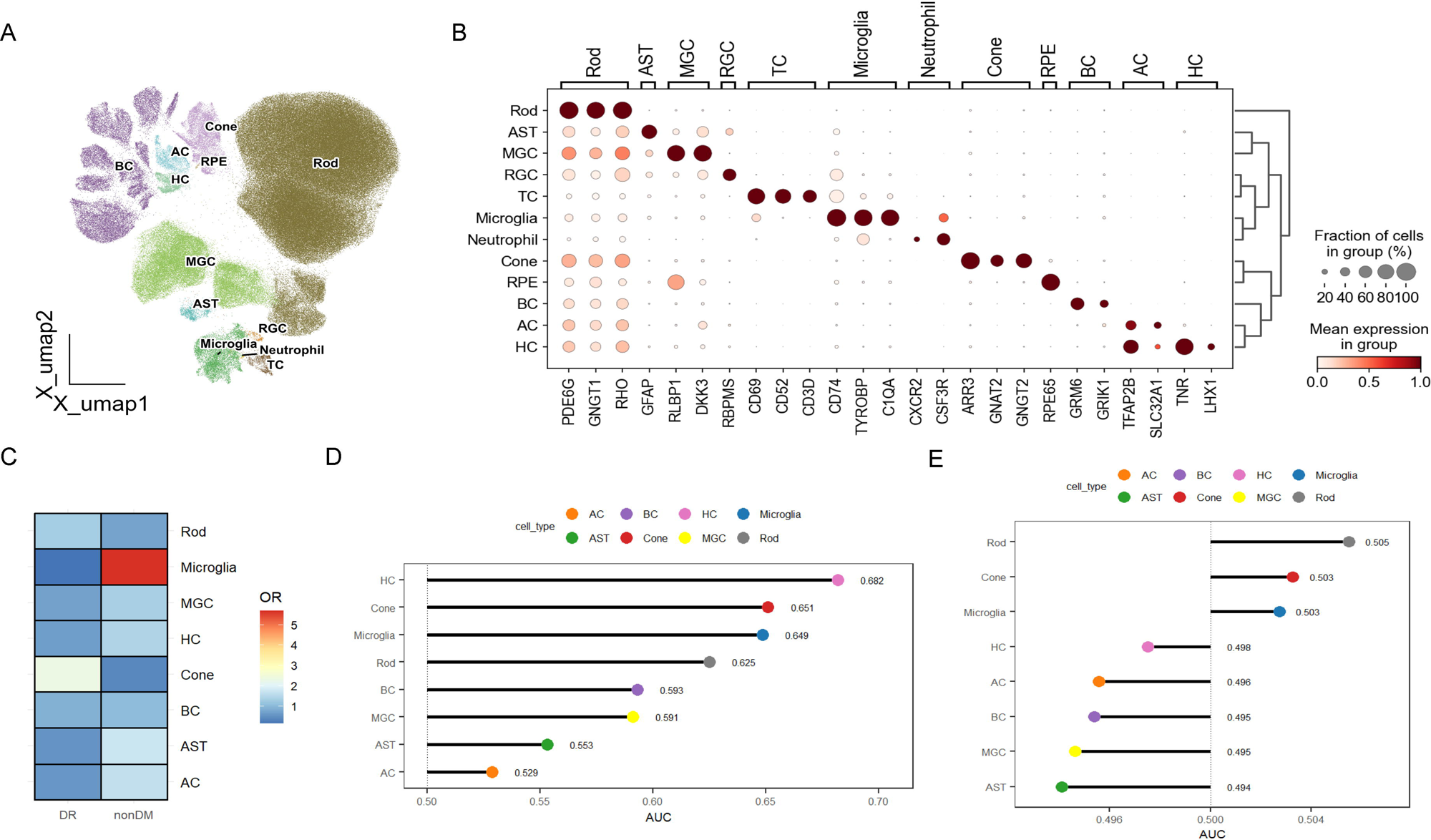
(A) UMAP visualization of major retinal cell types identified from scRNA-seq data. Cell types are color-coded and labeled. BC: Bipolar cell, MGC: Müller glia cell, TC: T cell, AC: Amacrine cell, RGC: Retinal ganglion cell, HC: Horizontal cell, RPE: Retinal pigment epithelium, AST: Astrocytes. (B) Dot plot of selected marker genes for major retinal cell types and T cells. Dot size represents the percentage of cells in each population expressing the gene, while color intensity indicates the normalized average expression level. (C) Odds ratio (OR) heatmap showing compositional enrichment of cell types between non-diabetic (nonDM) and diabetic retinopathy (DR) conditions. Within each condition column, warmer colors indicate OR>1 (enrichment in that column’s condition) and cooler colors indicate OR<1 (depletion). (D) Augur analysis ranks retinal cell populations by their predictive accuracy (AUC) in distinguishing nonDM and DR samples. Higher AUC scores correspond to greater changes in population frequency during disease progression. Each dot represents a specific cell type, color-coded by identity. (E) Label-permutation control for Augur. Randomizing condition labels yields AUC ≈ 0.5 across cell classes, confirming that the separability observed in (D) reflects genuine condition–expression structure rather than model artifact.

### Retinal C1QA+ myeloid cells resolve into homeostatic and inflammatory microglia plus a neutrophil cluster

To further characterize microglial heterogeneity, we sub-clustered the major *C1QA*-positive population, comprising 7,352 cells. This analysis resolved three subpopulations: homeostatic microglia (MG-Homeostatic), inflammatory microglia (MG-Inflammatory), and neutrophils (Figure 2A). Quality control metrics supported genuine biological heterogeneity (Figure 2B): MG-Inflammatory exhibited higher transcriptional complexity (nFeature_RNA, nCount_RNA), neutrophils the lowest, and all clusters showed uniformly low mitochondrial gene percentages (percent.mt), indicating high viability. Cluster-defining signatures from COSG (Figure 2C, 2D) revealed MG-Homeostatic enrichment for core microglial markers (*C1QA*, *C1QB*) and ribosomal protein genes (e.g., *RPL14*, *RPS4X*), whereas MG-Inflammatory displayed an activated profile with upregulation of immune-related genes (e.g., *DOCK4*, *CD83*, *KLF6*). The third cluster was identified as neutrophils by canonical markers (*FCGR3B*, *S100A12*, *CXCR2*) with minimal microglial gene expression. All three populations were observed across non-DM, DM, and DR retinas from multiple donors, indicating that clustering was not driven by condition or batch (Figure 2E, Table S2).

**Figure 2.**
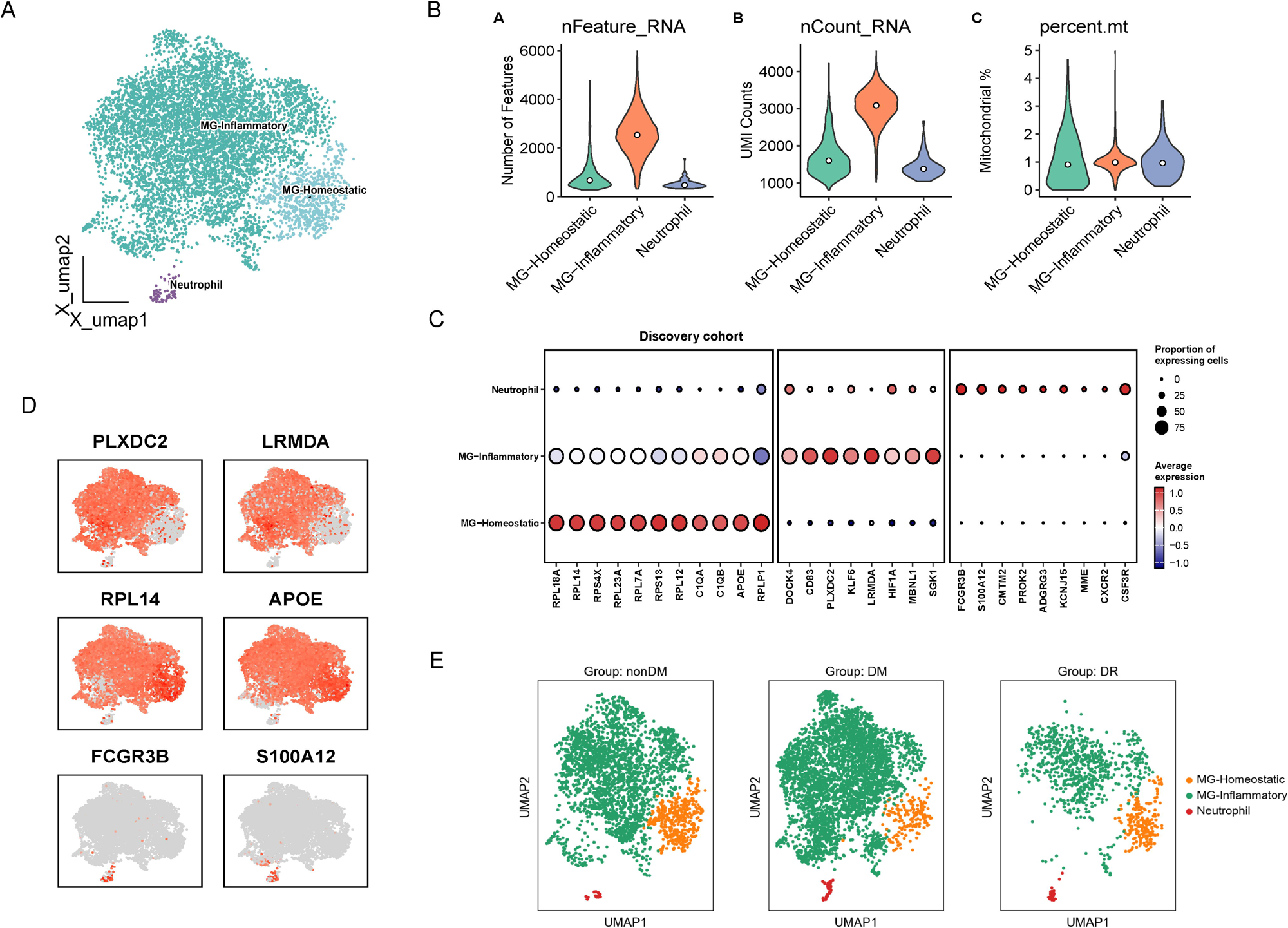
(A) UMAP visualization of C1QA+ myeloid cell subpopulation. Three distinct populations are labeled: MG-Homeostatic (homeostatic microglia), MG-Inflammatory (inflammatory microglia), and Neutrophil. (B) Violin plots showing quality control metrics across the three myeloid subpopulations. Panel A displays the number of detected genes (nFeature_RNA), Panel B shows the total RNA count (nCount_RNA), and Panel C indicates the percentage of mitochondrial genes (percent.mt). (C) Dot plot displaying cluster-defining marker genes identified by COSG across the discovery cohort. Dot size represents the proportion of cells expressing each gene within a population, while color intensity indicates the normalized average expression level. (D) Feature plots showing the spatial distribution of six representative marker genes on the UMAP. (E) UMAP projections comparing the distribution of C1QA+ myeloid cell subpopulation across nonDM (non-diabetic), DM (diabetic), and DR (diabetic retinopathy) conditions.

Comprehensive differential expression further substantiated these cell identities (Table S3). After rigorously excising ribosomal and rod-photoreceptor genes, pathway enrichment on the MG-Homeostatic versus MG-Inflammatory contrast delineated two non-overlapping programs (Figure 3A, 3B): MG-Homeostatic was enriched for mitochondrial oxidative phosphorylation, aerobic respiration, proton transmembrane transport, and ATP metabolic processes, whereas MG-Inflammatory was enriched for immune response–activating/regulating signaling, antigen processing and presentation of exogenous peptide antigen, macroautophagy, small GTPase–mediated signaling, endocytosis, lipid and atherosclerosis, osteoclast differentiation, and protein processing in the endoplasmic reticulum. irGSVA on immune hallmark gene sets (Figure 3C), while not excluding ribosomal/rod genes, recapitulated the same directional split across cell populations, strengthening the pathway-level separation between inflammatory and homeostatic programs; notably, neutrophils displayed a distinct immune-hallmark profile consistent with their canonical identity, providing orthogonal validation of all three populations. Importantly, homeostatic enrichment for oxidative phosphorylation and ATP metabolism persisted after removal of ribosomal and rod-photoreceptor genes and coincided with higher expression of antigen presentation/surveillance components (*CD74*, *HLA-DRA*/*DPA1*/*DPB1*, *TYROBP*, *AIF1*), arguing against a low-quality or degenerative state. Collectively, these pathway-level readouts from complementary methods indicate that inflammatory microglia engage coordinated cytokine and NF-κB/TLR-linked cascades, whereas homeostatic microglia maintain preserved bioenergetic tone alongside baseline immune surveillance functions.

**Figure 3.**
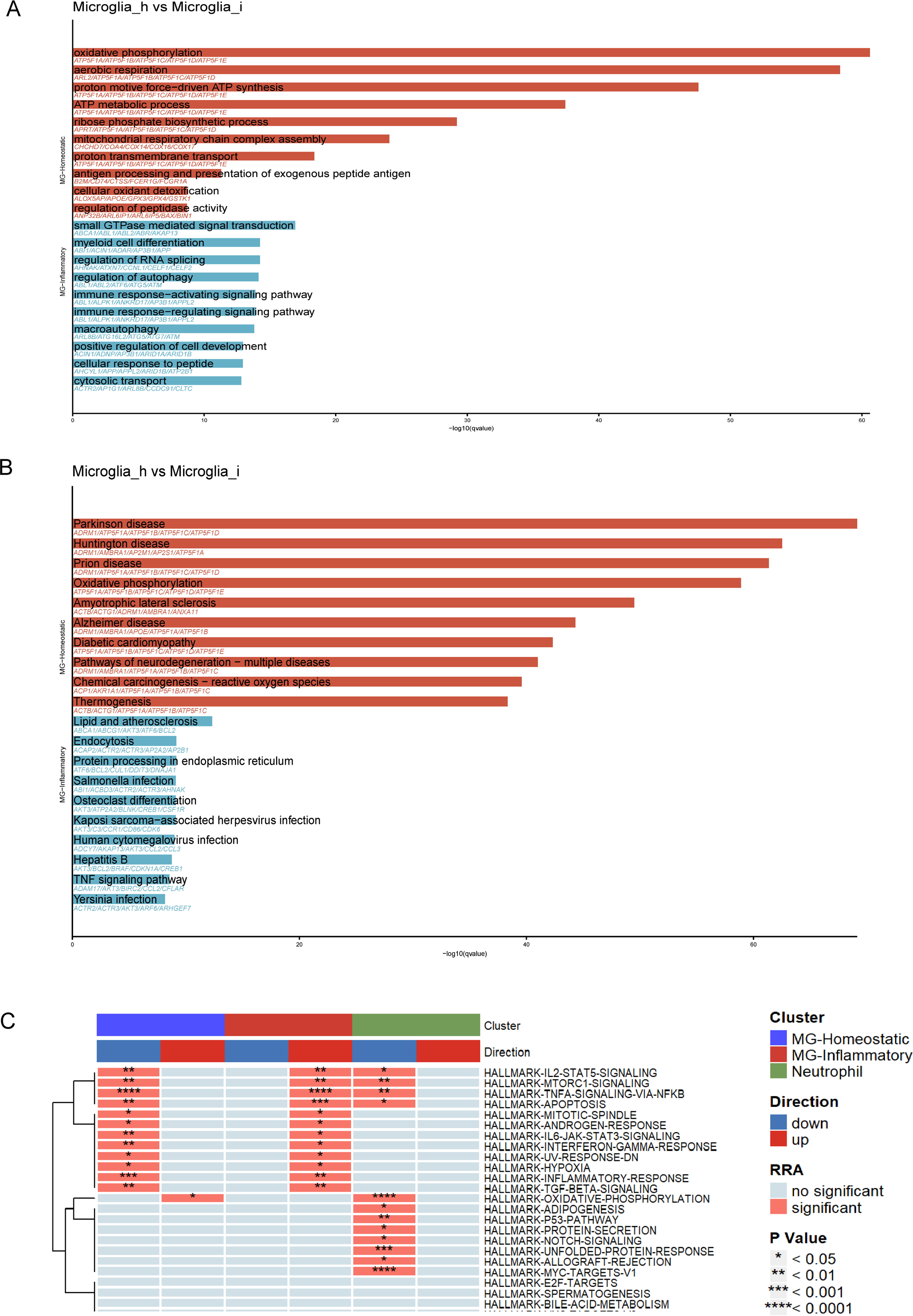
(A) Gene Ontology (GO) enrichment analysis showing significantly enriched biological processes between MG-Homeostatic (MG_h) and MG-Inflammatory (MG_i). Bar length represents -log10(adjusted p-value), with red bars indicating pathways enriched in MG-Homeostatic and blue bars showing pathways enriched in MG-Inflammatory. (B) KEGG pathway enrichment analysis comparing MG-Homeostatic (MG_h) versus MG-Inflammatory (MG_i). Bar length represents -log10(adjusted p-value), with red bars indicating pathways enriched in MG-Homeostatic and blue bars showing pathways enriched in MG-Inflammatory. (C) Heatmap of integrated gene set enrichment analysis (irGSEA) showing pathway activities across the three C1QA+ myeloid cell subpopulation. The top bar indicates cluster identity, and the second bar shows direction of regulation (blue: down, red: up). Significance levels are indicated by asterisks (* p < 0.05, ** p < 0.01, *** p < 0.001, **** p < 0.0001).

### Disease-stage dynamics of microglial subsets

Comparative analysis across disease states revealed distinct temporal activation patterns between the two microglial subsets (Figure 4). During early diabetic transition (DM vs. nonDM), MG-Homeostatic cells demonstrated functional stability with limited transcriptional changes, though notable upregulation of chemokines (*CCL2*, *CCL4*) and extracellular matrix remodeling genes (*FN1*) indicated early environmental sensing and low-grade inflammatory recruitment. Concurrently, transcriptional regulators (*NR4A2*) and neuroprotective factors (*CLU*) were enhanced, suggesting adaptive responses to hyperglycemic stress rather than overt activation. In contrast, MG-Inflammatory cells entered a highly active state characterized by significant upregulation of pro-inflammatory mediators (*IL6*), chemokines (*CCL2, CXCL2/3*), matrix-receptor and vascular regulatory molecules (*FN1, STAB1, SDC3, VAV3*), and copper oxidase (*CP*). Notably, classical lysosomal proteases (*CTSL*, *CTSZ*) were upregulated while *CTSH* was downregulated, indicating reconfiguration of phagocytic-proteolytic networks. The absence of significant *IL1A*/*IL1B* and *TNF* elevation confirmed this stage as “regulatory activation” rather than full inflammatory amplification.

**Figure 4.**
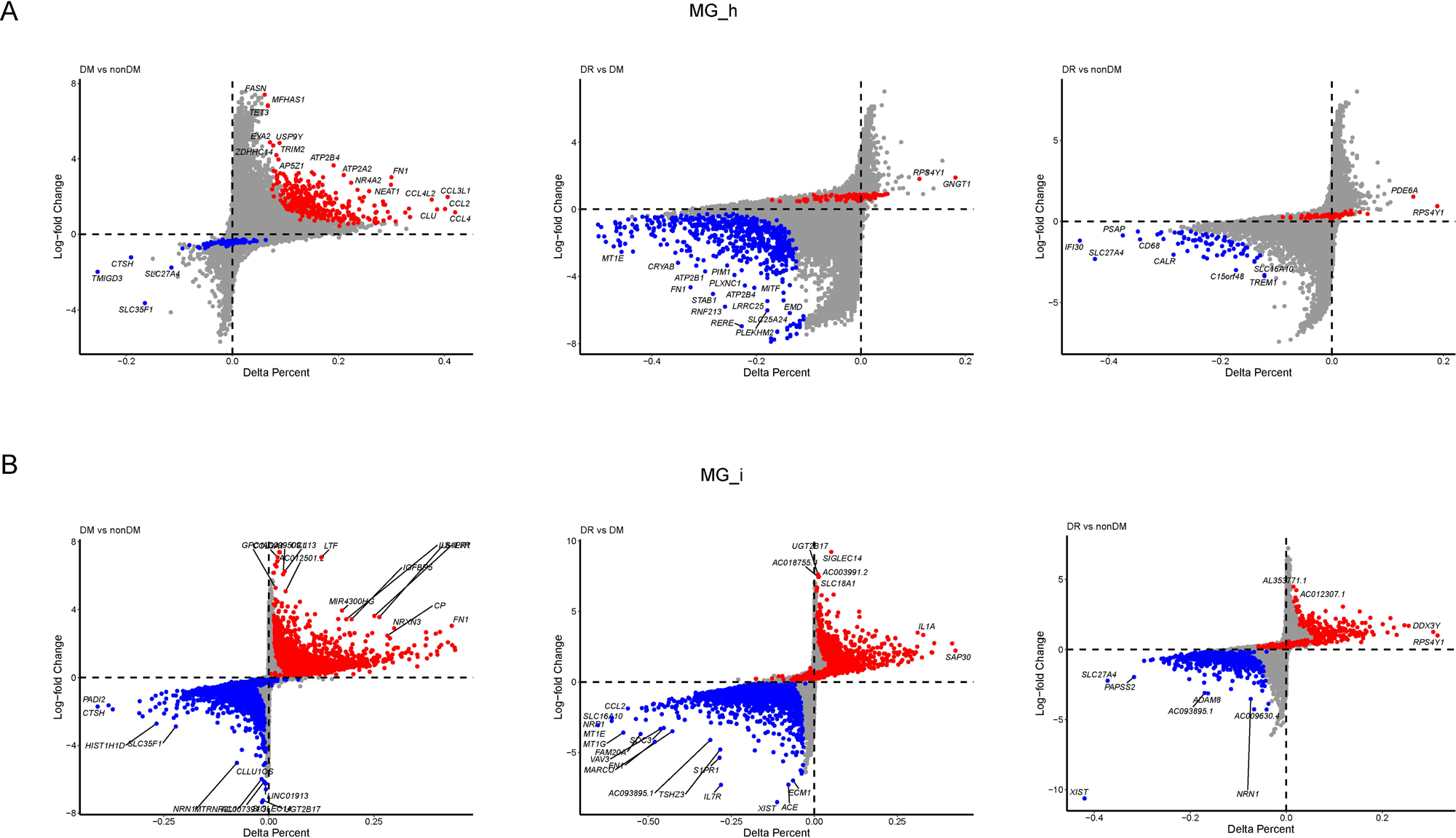
(A) Comparative differential gene expression analysis in MG-Homeostatic (MG_h) across disease states using diagonal scatter plots. Left panel: DM (diabetic without retinopathy) versus nonDM (non-diabetic control); middle panel: DR (diabetic retinopathy) versus DM; right panel: DR versus nonDM. For all plots, the x-axis represents delta percentage of gene expression (difference in the proportion of cells expressing a gene between the two conditions), and the y-axis shows the log2 fold change (LogFC) in expression levels. Red points indicate significantly upregulated genes (LogFC > 0.25, adjusted p-value < 0.05) in the first mentioned condition, while blue points indicate significantly downregulated genes (LogFC < -0.25, adjusted p-value < 0.05) in the first mentioned condition. Key marker genes are labeled. (B) Comparative differential gene expression analysis in MG-Inflammatory (MG_i) across disease states using diagonal scatter plots. Left panel: DM versus nonDM; middle panel: DR versus DM; right panel: DR versus nonDM. Axes and color coding follow the same convention as panel A.

Disease progression to retinopathy (DR vs. DM) deepened this functional divergence. MG-Homeostatic cells exhibited predominantly broad transcriptional downregulation in DR relative to DM, with upregulation largely confined to ribosomal protein genes (*RPLP1, RPS23, RPL41*); accordingly, these cells showed no evident induction of classical pro-inflammatory or angiogenic markers, consistent with a chronic, low-grade reactive state and diminished functional activity. Conversely, MG-Inflammatory cells demonstrated an intensified inflammatory program in DR, with *IL1A* and *IL1B* upregulated, and a reconfigured chemokine network evidenced by marked *CCL2* downregulation, alongside reduced expression of other chemokines (*CXCL1, CXCL2, CXCL3*). Vascular-related molecular remodeling was also evident in MG-Inflammatory cells, highlighted by *PDGFB* upregulation and *NRP1* downregulation, suggesting an enhanced capacity for vascular regulatory interactions. Direct comparison of DR versus nonDM further validated this divergent activation model, demonstrating that retinal microglia employ complementary rather than uniform activation strategies as disease progresses.

### Network-based analysis identifies microglial functional modules and disease-associated expression patterns

Gene co-expression analysis identified eleven discrete modules (M1-M11) with distinct molecular signatures (Figure 5A, Table S4). The correlation heatmap resolved two principal activation clusters: an inflammatory–stress cluster (M10, M4, M9) encompassing coordinated chemokine/cytokine, IL-1/immediate-early, and protein-stress programs; and a regulatory/metabolic–motility cluster (M5, M6, M8, M11) enriched for transcriptional control, lipid/metabolic genes, and remodeling/motility. M1 (ribosomal/translation) and M2 (rod-transcript signal) remained isolated; M2 is interpreted conservatively as photoreceptor carryover rather than a microglial program. M3 and M7 showed only moderate linkage to either cluster and were left ungrouped.

**Figure 5.**
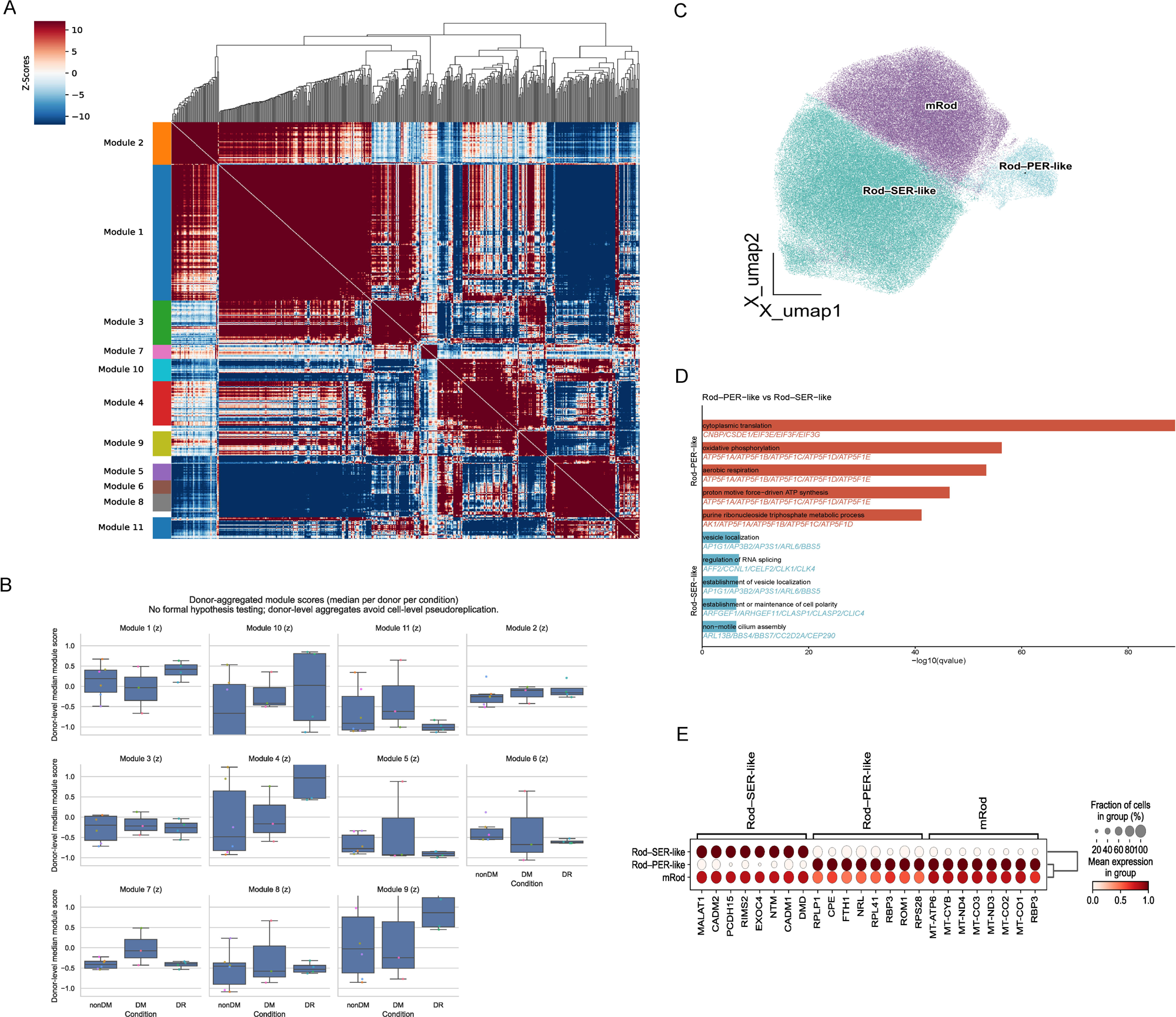
(A) Correlation heatmap showing relationships between identified Hotspot modules. Sidebar colors represent different module classifications. Color scale indicates correlation strength (red: positive correlation, blue: negative correlation) (B) Donor-aggregated module scores (median per donor per condition) across disease stages (nonDM: non-diabetic control, DM: diabetic without retinopathy, DR: diabetic retinopathy) for representative modules. (C) UMAP visualization of rod photoreceptor subclusters identified. Three functionally distinct rod subpopulations are resolved: Rod-SER-like (synaptic transmission-efficient rods with synaptic/adhesion program), Rod-PER-like (phototransduction-efficient rods with phototransduction/mitochondrial ETC/ATP program), and mRod (intermediate subtype with mixed gene-expression patterns). (D) Gene Ontology (GO) enrichment analysis comparing Rod-PER-like versus Rod-SER-like photoreceptors. Bar length represents -log10(adjusted p-value), with red bars indicating pathways enriched in Rod-PER-like and blue bars showing pathways enriched in Rod-SER-like. (E) Dot plot of selected marker genes for rod subtypes. Dot size represents the percentage of cells in each population expressing the gene, while color intensity indicates the normalized average expression level.

Consistent with this modular architecture, sample and donor level module scores (Figure 5B) showed that M1 was stable across nonDM, DM, and DR, with tightly clustered donor medians, indicating preserved baseline protein-synthesis capacity. In contrast, inflammatory–stress modules were higher in DR: M4 (IL1B with immediate-early TFs *JUN/FOS/EGR1/ATF3*), M10 (chemokines/cytokines *CCL3/CCL4/CXCL8/TNF*), and M9 (protein-stress/chaperones *HSPA1B/DNAJB1/HSPB1* and *DDIT4*) showed elevated scores in most DR donors. Distributions overlapped across groups and varied by donor, indicating heterogeneous activation. The DM group was generally intermediate/near baseline.

### Rod photoreceptor subtype persistence in DR disease stages

Previous studies have identified functionally distinct rod photoreceptor subtypes, phototransduction-efficient rods (PER) specialized for light detection and mitochondrial ATP synthesis, and synaptic transmission-efficient rods (SER) enriched in neurotransmission processes. Unsupervised reclustering resolved three rod groups: Rod–SER-like (synaptic/adhesion program), Rod–PER-like (phototransduction/mitochondrial ETC–ATP program), and an intermediate mRod with mixed gene-expression patterns (Figure 5C). Group-specific enrichment and differential-expression analyses aligned with previously reported PER/SER program (Figure 5D, 5E). MALAT1, which has been reported to decline with longer post-mortem intervals, ranked among the top differentially expressed transcripts between SER-like and PER-like groups. To assess potential source effects, we compared live and cadaver specimens and found that SER-like rods were more abundant in cadaver samples (overall mean 63.0% ± 10.9%) than in live samples (41.6% ± 21.2%), whereas PER-like rods were relatively higher in live samples (6.72% ± 7.30%) than in cadavers (3.51% ± 3.00%), indicating a retrieval-status bias in this cohort (Table S5). All three groups were detected across nonDM, DM, and DR, with inter-donor variability but no consistent stage-specific depletion.

### Rod subtypes and microglial states engage distinct signaling repertoires consistent with their transcriptional programs in nonDM condition

In nonDM conditions, rod cell subtypes exhibit distinct communication profiles (Figure S1A-C). PER-like rods display a secreted-factor bias, with prominent MIF-CD74 to inflammatory microglia and VEGFB-FLT1 to glia, suggesting metabolic/vascular support, while contributing little to rod–rod adhesion. In contrast, SER-like rods are dominated by synaptic/adhesive signaling and show reduced MIF/VEGF output compared to PER-like rods and show NRG1-ERBB4 signaling to interneurons and astrocytes, consistent with neuron-glia crosstalk. The mRod population exhibits a hybrid/transition profile, carrying PER-like ECM/vascular axes (VEGFB, VTN, PSAP) together with SER-like synaptic/trophic axes (NRG1, NOTCH), indicating a transitional rather than distinct identity. Overall, PER-like rods act as cytokine/growth-factor emitters with limited physical contact, whereas SER-like rods prioritize structural connectivity and synaptic support, aligning with the PER/SER transcriptional dichotomy. For retinal C1QA+ myeloid subpopulations, MG-Homeostatic engage in maintenance-oriented signaling, sending SPP1 and participate in MIF and C3 signaling (Table S6). MG-Inflammatory display an activated profile: they are the primary recipients of rod-derived MIF via CD74, and also take in TNF/TNFSF12, TGFβ1/2, ICAM1-β2-integrins, and extensive HLA-II-CD4 interactions. Neutrophils exhibit enriched chemotactic and adhesion signaling, consistent with their reactive characteristics.

### Remodeling of Cell Communication from nonDM to DR

In DR, intercellular communication shifts markedly compared to non-diabetic controls (Table S7, Figure S2). The pro-inflammatory MIF–CD74 axis remains active but skews toward the CXCR4 co-receptor branch in DR, while the CD44-dependent branch is relatively diminished. Inflammatory microglia (CD74-expressing) are the chief recipients of MIF in both states, with photoreceptors—especially the PER-like rods—as prominent sources; notably, DR boosts MIF signaling via CD74+CXCR4 at the expense of the CD74+CD44 pathway. By contrast, the neuroprotective PSAP–GPR37 interactions collapse in DR: robust microglia-to-Müller glia and rod-to-glia PSAP signals evident in non-diabetic (nonDM) conditions are almost absent in DR. Likewise, several trophic/metabolic support axes decline. Vitronectin–integrin signaling from rods virtually disappears in DR, as do NAMPT–INSR (visfatin to insulin receptor) interactions and FGF1–FGFR1 signaling that were detectable in the healthy retina. On the myeloid side, microglial outgoing signals become more limited. While osteopontin (SPP1) signaling is substantially reduced rather than intensified, it remains a relatively prominent inflammatory output from microglia as other signals also wane.

## Discussion

The advent of single-cell omics has significantly advanced our understanding of the pathological mechanisms underlying DR [9]. However, significant gaps persist, particularly in the absence of a comprehensive analysis of human DR retina samples. While animal models, including diabetic mice and non-human primates, have provided valuable insights [23–25], they fail to replicate the complexity of human DR pathogenesis due to species-specific differences. Recent efforts have partially addressed these gaps through single-cell analyses of human-derived tissues, such as PDR-fibrovascular membranes or blood mononuclear cells [26, 27]. However, the lack of studies involving human retina limits our ability to fully unravel the intricate cellular interactions and molecular mechanisms of DR in human.

To address this gap, we performed scRNA-seq on 20 human retinal samples covering three disease stages (nonDM, DM, and DR) from both surgical patients and donors. This comprehensive atlas spans all major retinal cell classes, including rod and cone photoreceptors, interneurons, Müller glia, vascular cells, and resident immune cells. In particular, the inclusion of microglia (resident immune cells known to mediate retinal inflammation [28, 29]) enables examination of their states across disease.

The molecular heterogeneity and functional dynamics of retinal microglia in DR are not fully understood due to insufficient comprehensive human retinal studies [28]. Traditionally described using the M1/M2 dichotomy, recent findings suggest that microglial activation spans a continuous spectrum rather than being defined by discrete phenotypes [28, 30]. Many studies that report “multiple microglial subpopulations” primarily reflect methodological advances that refine how putative groups are detected (e.g., deeper graph-based clustering, contrastive/representation learning, and improved batch correction) [31, 32]. Yet on low-dimensional visual embeddings, it often remains difficult to justify an additional, cleanly separable “third” cluster. In our dataset, we no longer find stable evidence for over two fully independent microglial “states” based on scRNA-seq human retina tissues, even though our atlas comprising 20 samples. Within the C1QA+ myeloid population, we resolved three subpopulations: MG-Homeostatic microglia, MG-Inflammatory microglia, and a neutrophil cluster. MG-Homeostatic cells expressed core microglial markers and ribosomal protein genes, whereas MG-Inflammatory cells upregulated immune-activation genes. Pathway analysis showed that MG-Homeostatic cells were enriched for oxidative phosphorylation and metabolic processes, while MG-Inflammatory cells were enriched in cytokine/NF-κB and antigen-presentation pathways. These findings reveal that retinal myeloid cells span a spectrum from quiescent energy-maintaining states to pro-inflammatory activation. Comparison across disease stages revealed distinct dynamics for these subsets. During early diabetes, MG-Homeostatic cells remained largely stable, aside from modest upregulation of chemokines and stress regulators, suggesting an adaptive response to hyperglycemia. In contrast, MG-Inflammatory cells became highly active in the DM stage, upregulating pro-inflammatory mediators, chemokines, and matrix/vascular regulators. As DR developed, MG-Homeostatic cells showed broad downregulation of gene expression with no new inflammatory markers, reflecting a suppressed profile, whereas MG-Inflammatory cells further intensified their inflammatory program. The identification of homeostatic and inflammatory microglial subsets, while initially appearing to support a binary framework, actually reveals a more sophisticated biological reality when integrated with our disease-stage dynamics and co-expression network analyses.

The molecular architecture underlying this continuum becomes apparent through our modular analysis, which reveals how microglial activation operates through coordinated recruitment of distinct functional programs rather than wholesale phenotypic switching. The inflammatory-stress axis (encompassing IL-1β/immediate-early, chemokine/cytokine, and protein-stress modules) progressively accumulates activity in DR while core biosynthetic machinery (translation module M1) remains remarkably stable across disease stages. This modular organization provides a mechanistic explanation for the observed divergent trajectories: homeostatic cells maintain their energetic backbone while selectively upregulating stress-response elements, whereas inflammatory cells layer additional pro-inflammatory programs onto this preserved foundation.

Moreover, our analysis confirms functionally distinct rod photoreceptor subtype populations persist consistently across the diabetes disease spectrum, with an additional intermediate mRod subtype displaying mixed transcriptional features. Crucially, these rod subtypes engage distinct communication repertoires that fundamentally shape retinal immune dynamics. PER-like rods function as primary secreted-factor emitters, prominently signaling through MIF-CD74 to inflammatory microglia and VEGFB-FLT1 to glial cells, while SER-like rods prioritize synaptic connectivity through NRG1-ERBB4 interactions with interneurons and astrocytes. This functional specialization becomes pathologically relevant in DR, where intercellular communication networks undergo systematic remodeling. Most significantly, the pro-inflammatory MIF-CD74 axis remains active but undergoes a critical qualitative shift toward CXCR4 co-receptor signaling in DR while CD44-mediated branches decline. This CD44 to CXCR4 transition reflects a shift from tissue-protective inflammation toward chronic, potentially tissue-damaging inflammatory signaling, as CXCR4 engagement typically promotes sustained inflammatory responses compared to the more regulatory CD44 pathway. Simultaneously, this inflammatory persistence occurs against a backdrop of collapsed neuroprotective networks. This coordination of maintained inflammatory signaling with lost homeostatic support reveals DR as fundamentally a disease of disrupted cellular cooperation, where specialized neuronal subtypes shift from tissue-protective to tissue-damaging communication patterns while losing their capacity to maintain retinal homeostasis.

While our study provides insights into DR progression, we acknowledge several limitations. The relatively small sample size may limit statistical power for certain comparisons. However, this limitation reflects the challenges in acquiring human retina samples, especially from DR patients. The human retina is exceptionally rare and difficult to obtain due to several factors: (1) the limited availability of donor tissue meeting specific disease criteria, (2) the critical requirement for minimal post-mortem intervals to maintain RNA quality, and (3) the ethical and logistical complexities of collecting samples from living donors during surgery. Despite these constraints, our study represents one of the largest single-cell analyses of human retina in DR research to date. The inclusion of both living donor and post-mortem specimens with careful processing time control helps mitigate some limitations while maintaining data quality. Future studies with larger sample sizes would be valuable to validate our findings, though the inherent challenges of human retinal tissue collection will likely continue to pose significant barriers to large-scale analyses.

## Statements and Declarations

### Data availability

The single-cell RNA sequencing data generated in this study have been deposited in Figshare with the DOI: 10.6084/m9.figshare.28228949.

### Authors’ Contributions

LY drafted the paper and performed the analysis. SL, YT, QP, TC contributed to data formatting and correction. YY, JL, YZ, QY, ZL, LC, GM, and RR provided comments on the paper. YS, and WM organized the project and provided comments. Collaborators from Queen’s University Belfast and Universiti Brunei Darussalam provided academic guidance and editorial suggestions only; they did not participate in data analysis, sample handling, or access to raw data. All substantive research activities were completed within Mainland China.

## Acknowledgment

This study was mainly funded by the Pioneer and Leading Goose R&D Program of Zhejiang Province 2023 with reference number 2023C04049 and Ningbo International Collaboration Program 2023 with reference number 2023H025.

## Corresponding authors

Correspondence to Dr. Weihua Meng and Dr. Yongqing Shao.

## Consent to Publish

All authors have consent for publication.

## Conflict of Interest

The authors have no conflicts of interest to disclose.

## Ethics approval

This study was approved by the Ethics Committee of the University of Nottingham Ningbo China; The Lihuili Hospital affiliated with Ningbo University; The Affiliated Ningbo Eye Hospital of Wenzhou Medical University.

---

**Figure S1.**
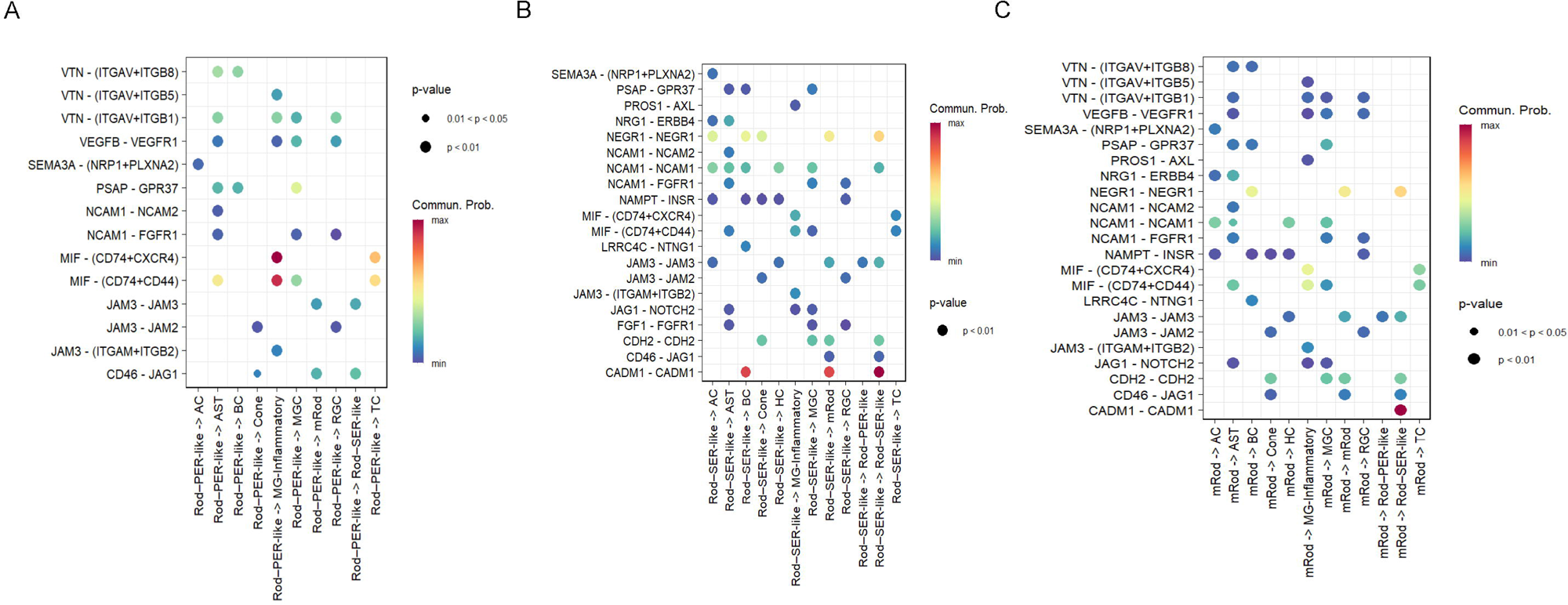
(A–C) Dot plots of sender-specific interactions inferred by CellChat, shown separately for Rod-PER-like (A), Rod-SER-like (B), and mRod (C) subtypes as senders toward the indicated receiver cell classes (x-axis). The y-axis lists selected ligand-receptor pairs highlighted in the Results. Color encodes CellChat communication probability (higher = stronger inferred interaction), and dot size encodes significance (larger dots: P < 0.01; smaller dots: 0.01 ≤ P < 0.05).

**Figure S2.**
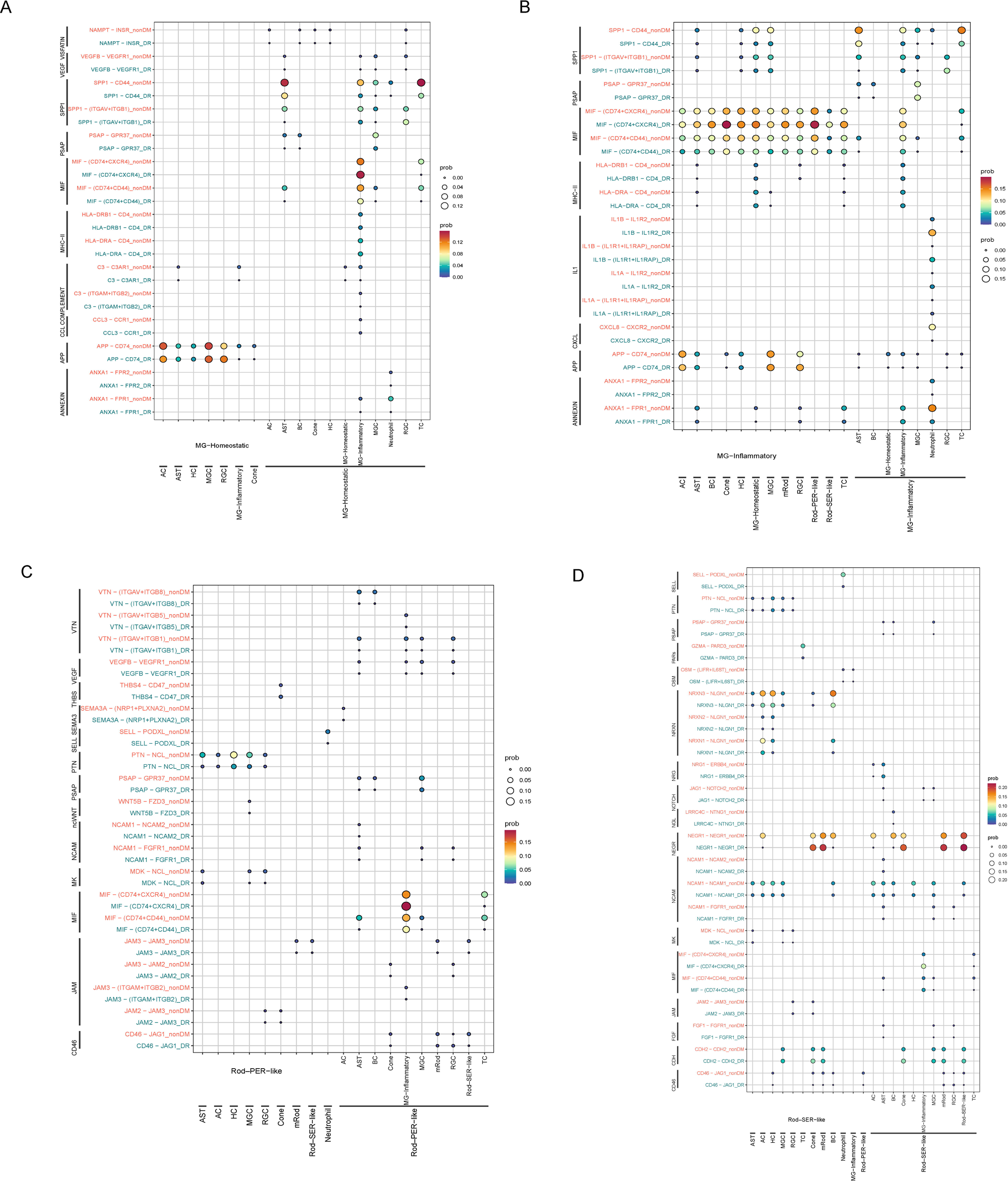
(A-D) Dot plots of both sender and receiver interactions inferred by CellChat, shown separately for MG-Homeostatic (homeostatic microglia) (A), MG-Inflammatory (inflammatory microglia) (B), Rod-PER-like (phototransduction-efficient rods) (C), and Rod-SER-like (synaptic transmission-efficient rods) (D). The y-axis lists selected ligand-receptor pairs, while the x-axis shows the interacting cell types as either senders (left sections) or receivers (right sections). Color encodes CellChat communication probability (orange/red = higher probability indicating stronger inferred interaction; blue = lower probability), and dot size encodes significance (larger dots: P < 0.01; smaller dots: 0.01 ≤ P < 0.05).

## Notes

### Competing Interest Statement

The authors have declared no competing interest.

### Summary of Updates

The analysis and results is updated, and the sample size increase from 13 samples to 20 samples in total.

